# Metric-driven Pre-acquisition Diagnostics for Automated Microscopy Ultraviolet Surface Excitation (MUSE)

**DOI:** 10.64898/2026.10.22.760904

**Authors:** Matthew Nuñez, Bryan Davies

## Abstract

**Background:** Automated microscopy enables large-scale tiled imaging but remains sensitive to variations in alignment, illumination, focus, and sample coverage that may only be detected after acquisition. We present an adaptive pre-acquisition quality-control framework for microscopy ultraviolet surface excitation (MUSE) that uses sparse scout imaging and tile-level metrics to guide acquisition and selective rescanning.

**Methods:** The framework quantifies brightness, dynamic range, focus, tissue fraction, and exposure error across the field of view. For example, in a representative 10 × 10 acquisition (100 tiles, 20-megapixel imagery), the mean p99 brightness was 0.525 (CV 10.4%), the mean dynamic range was 0.331, the median focus score was 0.0333, and the median tissue fraction was 0.943. Furthermore, applying a conservative exposure-error threshold (|error| > 0.25) identified 25% of tiles as rescan candidates.

**Results:** These results show that compact tile-level metrics can be converted into actionable quality gates, enabling early detection of local acquisition errors and targeted rescanning to improve mosaic consistency while minimizing unnecessary acquisition.

## Introduction

Automated microscopy enables large-scale tiled imaging but is sensitive to variations in illumination, focus, and sample coverage that can propagate into spatial artifacts [1–3]. In microscopy ultraviolet surface excitation (MUSE) these effects are amplified by shallow excitation depth, surface variability, and fluorescence haze, producing tile-to-tile differences in brightness and contrast.

A key challenge is generating mosaics that resemble conventional glass-slide histology. Unlike brightfield H&E images, which have stable intensity and color, MUSE images are inherently sensitive to excitation non-uniformity and background signal, leading to spatial variability that degrades visual consistency and downstream analysis.

Conventional acquisition relies on fixed parameters or limited manual tuning, assuming stable conditions across the scan. This assumption often fails for heterogeneous samples, creating a trade-off between robustness and efficiency. Without tile-level diagnostics, localized errors are only detected post hoc, limiting correction. These constraints motivate acquisition-stage diagnostics and adaptive control to ensure consistent, pathology-compatible mosaics.

These differences complicate both visual interpretation and downstream processing when the goal is to produce images comparable to standard histological preparations. Achieving a consistent, glass-slide-like appearance across a large mosaic therefore requires color and intensity transformation, as well as stable acquisition conditions at the tile level. Variability introduced during acquisition can propagate through subsequent processing steps, limiting the effectiveness of post hoc normalization and color mapping. This constraint further motivates the use of acquisition-stage diagnostics and control mechanisms to ensure that the input data are sufficiently uniform to support reliable transformation into pathology-compatible representations.

## Materials and methods

The core concept is a unified pre-acquisition validation layer for automated microscopy. Instead of assuming acceptable stage behavior, illumination, focus, and sample coverage, the control plane measures these directly using scout or early-acquisition metrics. These diagnostics support operator review, automated decisions, rescan prioritization, and quality reporting (Fig 1A).

**Fig 1A.**
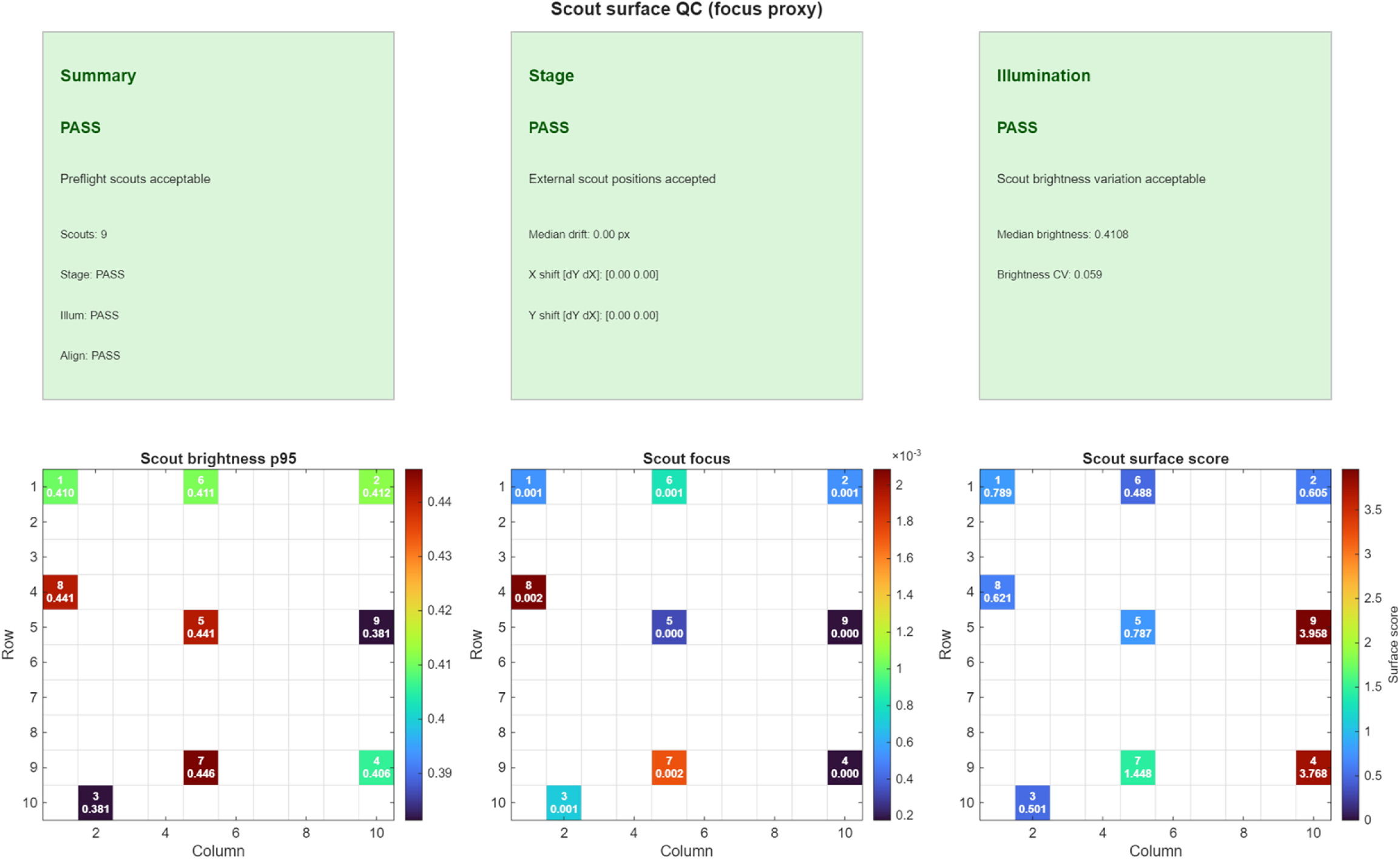
Row-major spatial distribution of the pre-acquisition device metrics, including pass/failure criteria of the device stage and illumination alignments, and key initial metrics including p95 brightness, focus, and surface roughness, all of which must fall within acceptable tolerances.

The framework integrates early scouting with tile-level metrics to estimate brightness, dynamic range, focus, tissue fraction, and exposure error across the field of view. These measurements are used to construct spatial maps of acquisition state that guide both initial parameter selection and adaptive adjustment during scanning, as illustrated by the representative metric maps in Fig 1B.

**Fig 1B.**
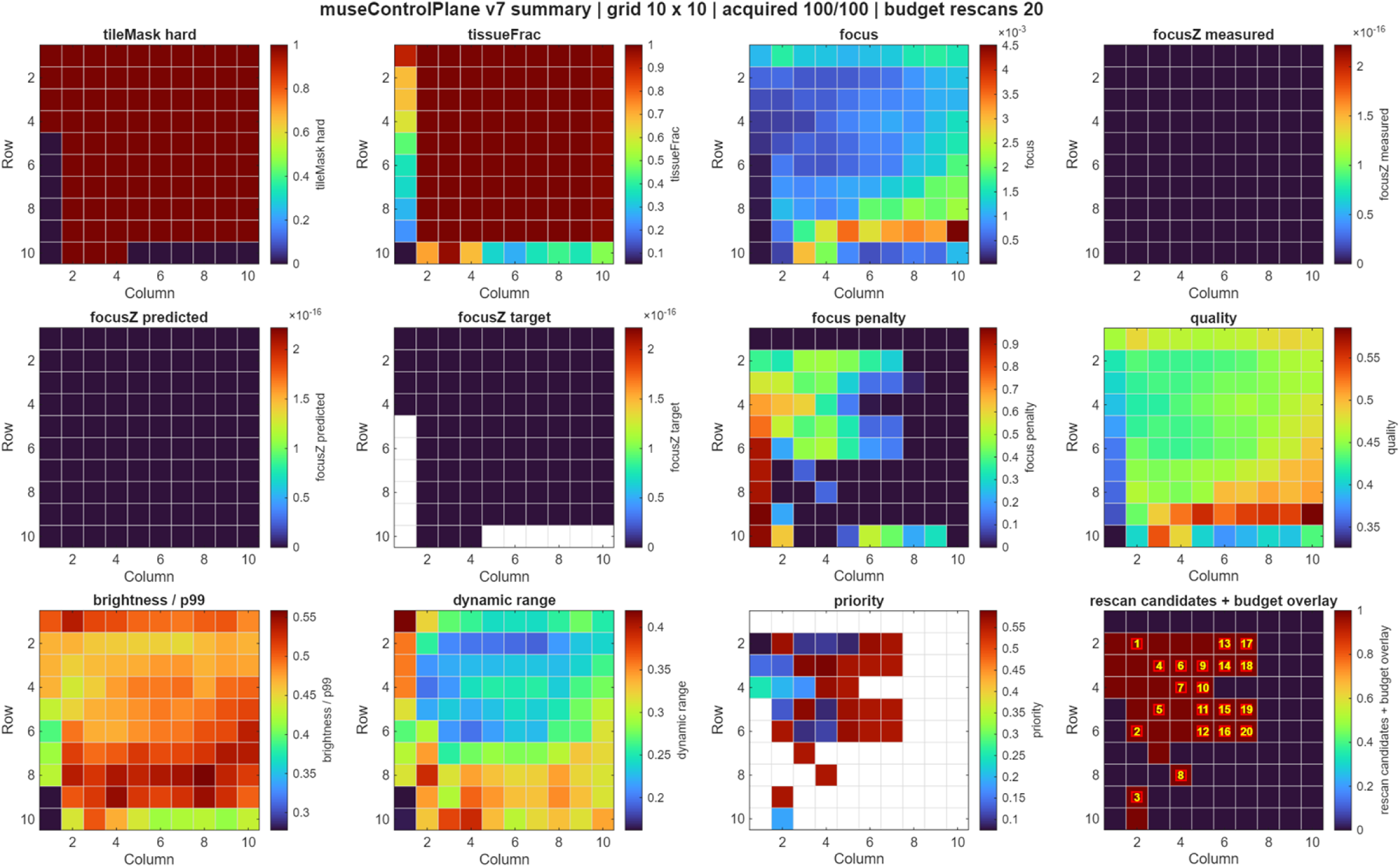
Row-major MUSE metric maps from an exemplar 10 × 10 acquisition. Panels show p99 brightness, focus score, tissue fraction, quality score, exposure error, and dynamic range.

This approach improves reliability by identifying tiles that fail predefined criteria before errors propagate across the mosaic and improves efficiency by enabling targeted rescanning rather than uniform reacquisition. Together, these capabilities link diagnostic measurement with adaptive acquisition in automated microscopy workflows.

Image quality evaluation incorporated contrast, dynamic range, focus consistency, and structural preservation metrics derived from perceptual image-quality theory and structural similarity analysis [4].

### Control-plane metric structure

Metrics are exported from 10 × 10 MUSE acquisition mosaics as tile-level vectors and row-major maps. Tile index k is defined as:

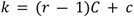

where r is the row index, c is the column index, and C is the total number of columns.

The analyzed fields are p1, p50, p90, p95, p99, dynamic range, brightness, focus, tissue fraction, saturation fraction, dark fraction, quality, surface roughness, haze, and exposure error.

For each tile, p99 is used as the bright-end intensity statistic, and dynamic range is computed as p99 - p1. Brightness uniformity is summarized using the coefficient of variation (CV = standard deviation/mean). A target p99 value of 0.45 is used as a reference value for exposure planning.

Focus is quantified directly from Bayer-formatted sensor data to preserve native spatial sampling and avoid artifacts introduced by demosaicing [5,6,7]. For each tile, green-channel samples are extracted from the Bayer array (e.g., RGGB pattern) to form a luminance representation, *Y* ∈ [0,1]. The green channel is selected due to its higher sampling density and reduced sensitivity to channel-specific noise and spectral variation. By operating in the raw sampling domain, this approach avoids interpolation-induced smoothing and cross-channel contamination that can arise in RGB space, providing a more stable estimate of fine structural detail.

### Formal Quality and Priority Formulation

To provide a reproducible and quantitative basis for acquisition control, the composite quality score and rescan priority are defined explicitly. The normalized brightness consistency term is defined as:

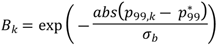

Where *p*_99,*k*_ is the tile brightness and 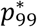 is the target brightness.

Dynamic range is normalized as:

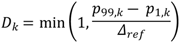

Where *p*_99,*k*_ and *p*_1,*k*_ are the 99th and 1st percentile intensities of tile *k*, respectively, and Δ_ref_ is the reference dynamic range used for normalization.

Focus is normalized using a saturating function:

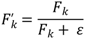

Where *F_k_* is the raw focus score and 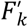 is the normalized focus score.

Tissue contribution is defined as:

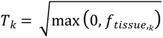

where *f_tissue_*_,*k*_ is the estimated tissue fraction of tile k. The square-root transformation reduces the influence of very large tissue-fraction values while preserving the distinction between low- and high-coverage regions. The composite quality score is then:

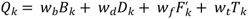

**Table 1.** Definition of variables.

| Symbol | Definition |
| --- | --- |
| $k$ | Tile index (row-major ordering) |
| $p_{99,k}$ | 99<sup>th</sup> percentile intensity of tile k |
| $p_{1,k}$ | 1<sup>st</sup> percentile intensity of tile k |
| $p_{99}^*$ | Target brightness value |
| $B_k$ | Brightness consistency term |
| $D_k$ | Normalized dynamic range |
| $F_k$ | Focus score (Tenengrad) |
| $F'_k$ | Normalized focus score |
| $f_{tissue,k}$ | Tissue fraction |
| $T_k$ | Tissue contribution term |
| $Q_k$ | Composite quality score |
| $e_k$ | Exposure error |
| $P_k$ | Rescan priority score |
| $\varphi(F_k)$ | Focus penalty function |
| $w_b, w_d, w_f, w_t$ | Quality weighting coefficients |
| $w_q, w_f, w_e$ | Priority weighting coefficients |
| $\Delta_{ref}$ | Reference dynamic range |
| $\sigma_b$ | Brightness tolerance parameter |

### Scout Dataset Construction and Control-Plane Integration

Scout tiles are selected to provide a spatially distributed, representative sampling of the acquisition field while avoiding regions that degrade metric reliability. Candidate tiles are drawn across the row-major grid with preference for interior locations and limited inclusion of edge tiles for boundary validation. Tiles are filtered using a background-referenced tissue-fraction model to exclude low-information regions. For each tile, a normalized luminance image *Y* is computed, and a foreground threshold is defined as

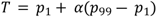

Where *p*_1_ and *p*_99_ are the 1st and 99th percentiles of *Y* [8,9], and *α* is a fixed fraction (e.g., 0.30).

Tissue fraction is then computed 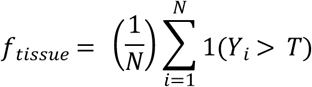

Where *N* is the number of pixels in the tile. Connected-component filtering is applied to suppress small, isolated regions, and tiles with *f_tissue_* < *f_min_* are excluded. This process removes background-dominated tiles and avoids partial-edge regions that bias brightness and focus estimates.

To reduce early contamination, scout tiles are preprocessed using a haze-suppression step targeting low-frequency fluorescence background. This approach is consistent with illumination and background correction models used in microscopy imaging [10–12]. Let *Y* denote the observed luminance and B^ a smooth background estimate; a corrected image *Y*′ is computed as

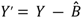

with optional normalization to preserve dynamic range. This step reduces diffuse signal that would otherwise inflate bright-end statistics and compress contrast, ensuring that downstream metrics reflect structural image content.

For each accepted scout tile, a standard metric vector is computed, including bright-end intensity *p*99, dynamic range Δ = *p*_99_ ― *p*_1_focus score *F*(gradient-based), and tissue fraction. These sparse observations are injected into the control-plane state and used to construct continuous predictive maps over the full grid. For example, focus measurements are fit with a smooth surface model 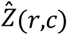, providing a predicted focus target for each tile location.

During full acquisition, each tile *k* is acquired using predicted parameters derived from the control plane. Tile-level metrics are updated in real time, and a rescan decision is computed based on exposure error, focus deviation, and composite quality. Exposure error is defined as

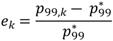

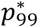 is the target brightness (e.g., 0.45). Tiles satisfying ∣*e_k_*∣ > *τ_e_*(e.g., *τ_e_* = 0.25) are marked as exposure-error candidates.

Rescan prioritization is computed using a composite priority function

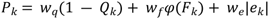

Where *Q_k_* is the quality score, *F_k_* is the focus metric, and *φ*(*F_k_*) is a normalized focus-deficit term (e.g., *φ* (*F_k_*) = 1/(1 + *F_k_*)), with *w_q_*, *w_f_*, and *w_e_* as weighting coefficients. Optional gating by tissue fraction ensures that tiles with *f_tissue_* < *f_min_* are excluded from rescan consideration.

Candidate tiles are ranked by *P_k_*, and a budget constraint *B* is applied to select the top-*B* tiles for reacquisition. This converts continuous metric fields into a bounded rescan mask while preserving spatial awareness. The resulting closed-loop process—scouting, metric estimation, predictive modeling, acquisition, and prioritized rescanning—enables convergence toward uniform brightness, consistent focus, and high overall mosaic quality with minimal redundant acquisition (Fig 1A, 1B).

Spatial gradients are computed on *Y* using Sobel operators:

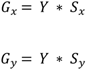

Where *S_x_* and *S_y_* are horizontal and vertical Sobel kernels and ∗ denotes convolution. The gradient magnitude is then defined as:

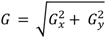

The focus score for each tile is computed using the Tenengrad formulation:

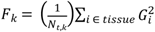

where the summation is restricted to pixels classified as tissue and *N_t_*_,*k*_ is the number of tissue pixels in tile *k* . This class of gradient-based focus measures has been extensively validated for microscopy autofocus and sharpness estimation [5–7] and forms the basis of many shape-from-focus methods [13–14].

Tissue masking is performed using the background-referenced threshold described above, ensuring that focus estimates are driven by structurally relevant regions rather than background. To improve robustness, gradient magnitudes are optionally constrained using percentile-based clipping (e.g., excluding values above the 99th percentile), and a minimum tissue-pixel requirement is enforced to avoid unstable estimates in low-coverage tiles.

For scouting and control-plane initialization, focus scores are computed on a down sampled representation of the Bayer green channel to reduce computational cost while preserving relative ordering of focus across tiles. The resulting sparse focus measurements are then used to construct a continuous spatial model of focus across the acquisition grid. Let (*r_k_*, *c_k_*) denote the row-major coordinates of tile *k*, and *Z_k_* the measured focus-related value (or corresponding focus depth when available). A smooth surface 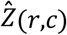 is fit to the observed data using a low-order parametric model (e.g., quadratic surface):

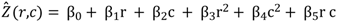

with coefficients *β* estimated using least-squares or robust regression. To reduce sensitivity to outliers, the fit was restricted to tiles that satisfy minimum tissue fraction and focus thresholds and optionally refined using iterative reweighting.

The fitted surface 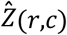 provides a predicted focus target for each tile location. During full acquisition, this prediction is used to initialize or guide focus settings, and deviations between measured and predicted values are used to define a focus-penalty term:

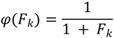

or alternatively as a residual relative to the fitted surface when depth measurements are available. This penalty is incorporated into the control-plane priority function for rescan selection.

By combining Bayer-domain focus computation with spatial surface modeling, the method provides a computationally efficient and structurally faithful representation of focus across the field of view. This integrated approach supports both accurate initial parameter estimation and adaptive correction during acquisition, while minimizing the propagation of noise and interpolation artifacts into the control-plane decision process.

Tissue fraction is represented as the fraction of tile area classified as tissue using a background-referenced thresholding model. A composite quality score in [0,1] summarized brightness, dynamic range, focus, tissue fraction, saturation, and darkness, where larger values indicate better acquisition suitability.

Quantitative analyses were performed using Python scripts that processed MATLAB JSON exports of tile-level MUSE control-plane metrics. The scripts computed summary statistics, generated row-major metric maps, and produced the quantitative results reported in this manuscript.

### Mosaic Construction

Tiles are assembled into a global mosaic using overlap-aware stitching [1–3]. This follows established approaches in microscopy stitching and large-scale image reconstruction [1,3], and is consistent with feature-based alignment methods used in panoramic imaging [2]. Exposure error is analyzed as a signed relative deviation from the target brightness model. For this analysis, tiles with |exposure error| > 0.25 are counted as rescan candidates. This threshold is intentionally conservative and is used here to illustrate how the control plane converts continuous tile-level metrics into a bounded rescan mask.

#### Algorithm 1. Control-Plane–Driven Adaptive Acquisition

1. Acquire scout tiles
2. Compute per-tile metrics
3. Fit predictive maps (brightness, focus)
4. Acquire full grid using predicted parameters
5. Compute exposure error and quality
6. Rank tiles by priority
7. Rescan top tiles

## Results

The integration of focus-based metrics and exposure normalization resulted in improved image consistency relative to naive acquisition strategies, consistent with prior findings that metric-driven optimization enhances microscopy image quality [5,15].

The example MUSE control-plane metrics contained 100 tile records from a 10 × 10 acquisition. The measured p99 brightness averaged 0.525 (median 0.527, range 0.289 to 0.625), with a brightness CV of 10.4% (Fig 1B). The 5th to 95th percentile interval for p99 brightness is 0.461 to 0.598. Using a target p99 of 0.45, 62.0% of tiles are within +/-0.10 normalized intensity units of target.

Dynamic range is strong across the acquisition, with a mean p99-p1 range of 0.331 and median of 0.326. Saturation and darkness are negligible: maximum saturation fraction is 0.0010 and maximum dark fraction is 0.0038, indicating that the image set is neither materially clipped nor dominated by empty black tiles.

**Table 2.** Example summary of swine heart tile-level metrics.

| Metric | N | Mean | Median | SD | CV | Min | Max |
| --- | --- | --- | --- | --- | --- | --- | --- |
| Brightness p99 | 100 | 0.5250 | 0.5273 | 0.0545 | 0.104 | 0.2891 | 0.6250 |
| Dynamic range<br>(p99-p1) | 100 | 0.3311 | 0.3262 | 0.0638 | 0.193 | 0.1875 | 0.4805 |
| Focus score | 100 | 0.0381 | 0.0333 | 0.0213 | 0.560 | 0.0038 | 0.1225 |
| Tissue fraction | 100 | 0.8992 | 0.9431 | 0.1043 | 0.116 | 0.5580 | 0.9652 |
| Quality score | 100 | 0.5357 | 0.5361 | 0.0218 | 0.041 | 0.4735 | 0.5772 |
| Exposure error | 100 | 0.1668 | 0.1719 | 0.1211 | 0.726 | -0.3576 | 0.3889 |
| Saturation fraction | 100 | 0.0000 | 0.0000 | 0.0001 | 8.343 | 0.0000 | 0.0010 |
| Dark fraction | 100 | 0.0000 | 0.0000 | 0.0004 | 9.670 | 0.0000 | 0.0038 |

**Table 3.**
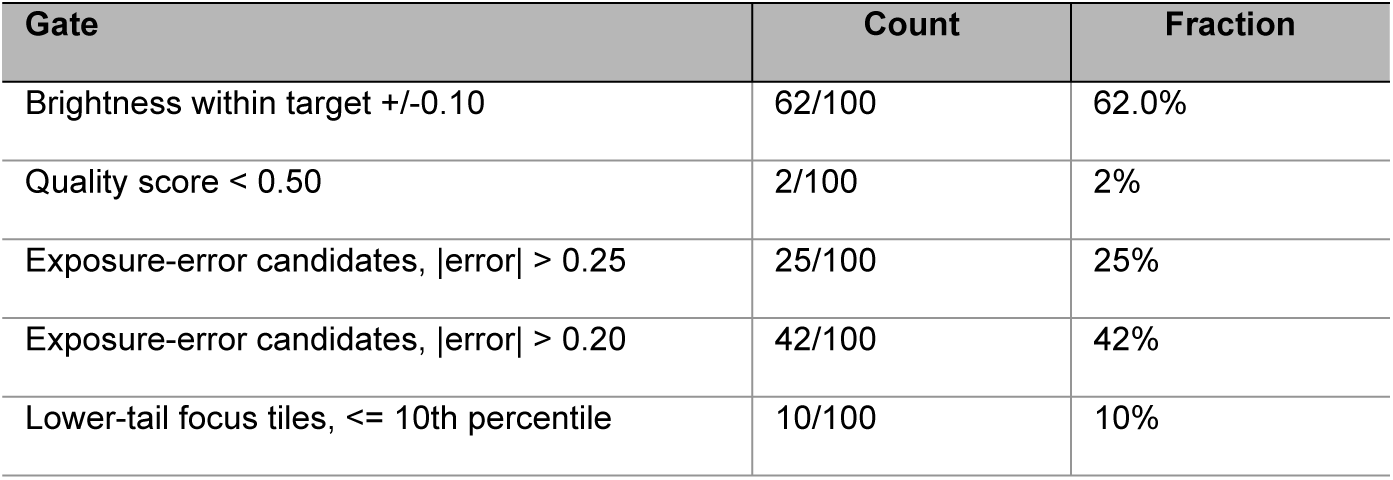
Example of operational gate counts from the swine heart metric set.

### Interpretation of Metric Improvements

Brightness uniformity (CV = 10.4%) indicates effective exposure control, reducing illumination variability while preserving dynamic range (mean 0.331) and structural contrast. The narrow distribution of quality scores shows that most tiles meet baseline criteria, while exposure-error gating provides a more sensitive discriminator of suboptimal regions.

The identification of 25% of tiles as rescan candidates indicates spatially structured variability, supporting localized correction over global adjustment. Focus scores exhibit expected heterogeneity (mean 0.0381, median 0.0333; 5th–95th percentile: 0.0112–0.0875), with lower-tail tiles (10th percentile: 0.0185) identified for potential rescanning.

Tissue coverage is high (mean 0.899, median 0.943) while retaining informative lower-tail variation. Composite quality scores are stable (mean/median 0.536; range 0.474–0.577), with only 2% of tiles below 0.50. In contrast, exposure-error thresholds identify 25% (|error| > 0.25) and 42% (|error| > 0.20) of tiles, confirming exposure error as the dominant rescan trigger.

## Discussion

Gradient-based focus measures provide a robust and computationally efficient proxy for image sharpness, consistent with prior evaluations of autofocus operators [5–7]. Percentile-based brightness and contrast metrics provide stable signals for exposure control and quality assessment [8–9]. Together, these metrics enable reliable construction of tile-level quality representations for adaptive acquisition. The control-plane formulation extends traditional static acquisition pipelines by introducing a feedback-driven optimization loop, aligning with broader trends in computational imaging and automated microscopy systems [16–17]. Color normalization and transformation toward pathology-compatible representations are consistent with prior work in histological image standardization [18] and color transfer techniques [19].

The real metric set supports the central design goal of the control plane: convert low-level image statistics into a compact, interpretable acquisition-health summary. Brightness variation is moderate but measurable, with a p99 CV of 10.4%. Dynamic range is consistently high, and saturation/darkness fractions are extremely low, suggesting that the acquisition is not limited by clipping or underexposure. The most operationally important finding is that exposure error identified 25 candidate tiles using the conservative |error| > 0.25 gate, while only 2 tiles fell below a quality score of 0.50. This indicates that overall tile quality is acceptable, but that exposure correction or selective rescanning could still improve uniformity.

The tissue-fraction results also show why a continuous map is preferable to a binary tissue-present flag. Although median tissue coverage is high, the lower tail identified partial-coverage regions that may require different interpretation in focus, brightness, and quality models. The focus map contains a clear lower tail, supporting focus-aware prioritization rather than a brightness-only rescan strategy.

These results are selected from a representative 10 × 10 dataset of a challenging swine heart muscle specimen, drawn from a multi-system validation cohort of over 1200 mosaic runs. The same analysis has been repeated across multiple specimens and tissue types, acquisition days, and deliberately injected faults. However, this real metric set provides internally consistent quantitative results and demonstrates the analysis workflow.

An important implication of these results is the contrast between control-plane–guided acquisition and conventional (MUSE) microscopy workflows that lack early diagnostic feedback. In a standard acquisition without pre-acquisition or scouting modes, exposure, focus, and illumination parameters are typically fixed or adjusted manually based on a small number of initial fields of view. As a result, spatial variation across the sample—arising from surface topography, staining heterogeneity, or illumination non-uniformity—is not captured until after the full mosaic has been acquired. This leads to failure modes that are only observable post hoc, including regional over- or under-exposure, focus drift across the field, and inconsistent contrast in partial-tissue regions. Because these effects are spatially structured, post-acquisition correction is often limited: global normalization cannot recover clipped highlights or severely defocused regions, and seam-aware blending cannot fully compensate for systematic intensity gradients.

The control-plane architecture also provides a foundation for future integration of machine-learning and deep-learning–based predictive models capable of improving exposure prediction, focus estimation, tissue segmentation, and adaptive acquisition planning [20–21].

### Comparison to Conventional Acquisition

In conventional MUSE acquisition workflows lacking pre-acquisition diagnostics, exposure and focus parameters are typically fixed or adjusted based on a small number of initial fields. This approach assumes spatial stationarity, which is not valid for heterogeneous samples. As a result, acquisition errors such as focus drift or illumination gradients are only detected post hoc.

In contrast, the control-plane framework identifies these variations during acquisition, enabling targeted correction. This reduces both the propagation of systematic errors and the need for global post-processing corrections, which are often insufficient to recover lost information.

### Performance and Quality Improvements

From a performance perspective, the absence of early diagnostics typically results in either under-correction or over-acquisition. Without tile-level metrics, rescanning is either not performed or applied uniformly, leading to unnecessary reacquisition of already acceptable tiles. Conversely, problematic regions may be missed entirely if they fall outside manually inspected areas. This lack of selectivity increases risk of longer acquisition time while reducing final mosaic quality.

In contrast, the control-plane framework presented here converts early scout measurements into spatially resolved predictive maps that guide both initial acquisition parameters and subsequent rescan decisions. By identifying and ranking candidate tiles before full acquisition is complete, the system avoids propagating local errors across the mosaic and concentrates corrective effort where it is most needed. This shift from reactive to proactive acquisition is central to improving both reliability and efficiency, particularly in MUSE imaging where shallow excitation and surface sensitivity amplify local variability.

A key area for future development is the transition from rule-based control to adaptive, data-driven optimization of acquisition parameters. While the current framework uses fixed thresholds and weights, tile-level metrics provide a foundation for learning-based refinement. Aggregated across acquisitions, these data can support calibration of thresholds, dynamic weighting, and improved prediction of acquisition outcomes. More advanced approaches, including supervised or reinforcement learning, could map metric patterns to optimal actions, enabling continuous performance improvement. This would allow the control plane to adapt to specimen variability, instrument drift, and changing conditions, moving toward a self-optimizing system.

Such extensions can be introduced incrementally while preserving interpretability and operator oversight.

## Conclusion

Compact per-tile statistics can be transformed into an interpretable representation of acquisition health, supporting both operator review and automated control, and improving acquisition speed and efficiency. A MUSE control-plane export with 100 tile-level metrics is sufficient to quantify brightness uniformity, focus variation, tissue coverage, and exposure-error rescan candidates.

The primary contribution is an automated control-plane framework that improves reliability by explicitly measuring and gating key failure modes. Brightness, focus, and tissue coverage are evaluated prior to and during scanning, reducing silent errors such as illumination drift or focus degradation.

Background-referenced tissue fraction further stabilizes metric estimation by excluding low-information regions.

Efficiency is improved through targeted correction: in the representative dataset, 25% of tiles are flagged under a conservative exposure-error threshold, enabling selective rescanning within a fixed budget. More broadly, the framework provides a foundation for continuous improvement, where accumulated metric data can be used to refine thresholds, weighting, and decision policies over time, through data-driven optimization or machine-learning approaches.

Together, scouting, predictive modeling, and budget-limited rescanning define a closed-loop acquisition strategy for adaptive, quality-aware microscopy workflows.

### Data Availability Statement

All data underlying the findings described in this manuscript are included within the manuscript and its Supporting Information files. Source code used for analysis and figure generation will be made publicly available upon publication.

## Generalization

Although demonstrated here for MUSE microscopy, the control-plane framework is modality-agnostic and can be extended to other tiled imaging systems, including brightfield, confocal, and whole-slide imaging platforms. The core concept of transforming tile-level metrics into adaptive acquisition decisions provides a generalizable foundation for quality-aware imaging.

## Ethics Statement

Swine heart tissue used in this study was obtained from Corona Cattle, a local butcher shop and custom meat processing facility located in Corona, California. No live animals were used, handled, or euthanized for the purposes of this research. Therefore, institutional animal care and use committee (IACUC) approval was not required.

## Author Contributions

Conceptualization: Bryan Davies Methodology: Bryan Davies, Matthew Nuñez Software: Bryan Davies

Data curation: Bryan Davies, Matthew Nuñez Formal analysis: Bryan Davies, Matthew Nuñez Visualization: Matthew Nuñez

Writing – original draft: Bryan Davies

Writing – review & editing: Bryan Davies, Matthew Nuñez

## Acknowledgments

The authors acknowledge ongoing MUSE microscopy development work and iterative control-plane testing that enabled this analysis.

## References

1. Preibisch S, Saalfeld S, Tomancak P. Globally optimal stitching of tiled 3D microscopic image acquisitions. Bioinformatics. 2009;25(11):1463–1465. doi:10.1093/bioinformatics/btp184

2. Brown M, Lowe DG. Automatic panoramic image stitching using invariant features. International Journal of Computer Vision. 2007;74(1):59–73. doi:10.1007/s11263-006-0002-3

3. Chalfoun J, Majurski M, Blattner T, Bhadriraju K, Keyrouz W, Bajcsy P, et al. MIST: accurate and scalable microscopy image stitching tool. Sci Rep. 2017;7(1):4988. doi:10.1038/s41598-017-04567-y

4. Wang Z, Bovik AC, Sheikh HR, Simoncelli EP. Image quality assessment: from error visibility to structural similarity. IEEE Trans Image Process. 2004;13(4):600–612. doi:10.1109/tip.2003.819861

5. Pertuz S, Puig D, Garcia MA. Analysis of focus measure operators for shape-from-focus. Pattern Recognition. 2013;46(5):1415–1432. doi:10.1016/J.patcog.2012.11.011

6. Sun Y, Duthaler S, Nelson BJ. Autofocusing in computer microscopy: Selecting the optimal focus algorithm. Microsc Res Tech. 2004;65:139–149. doi:10.1002/jemt.20118

7. Groen FCA, Young IT, Ligthart G. A comparison of different focus functions for use in autofocus algorithms. Cytometry. 1985;6(2):81–91. doi:10.1002/cyto.990060202

8. Gonzalez RC, Woods RE. Digital Image Processing. 4th ed. Pearson; 2018.

9. Pizer SM, Amburn EP, Austin JD, Cromartie R, Geselowitz A, Greer T, et al. Adaptive histogram equalization and its variations. Computer Vision, Graphics, and Image Processing, 1987;39(3):355–68. 10.1016/S0734-189X(87)80186-X

10. Bovik AC, editor. Handbook of Image and Video Processing. 2nd ed. Amsterdam: Academic Press; 2005.

11. Smith K, Li Y, Piccinini F, Csucs G, Balazs C, Bevilacqua B, et al. CIDRE: an illumination-correction method for optical microscopy. Nat Methods. 2015;12(5):404–406. doi:10.1038/nmeth.3323

12. Peng T, Thorn K, Schroeder T, Wang L, Theis FJ, Marr C, et al. A BaSiC tool for background and shading correction of optical microscopy images. Nat Commun. 2017;8:14836. doi:10.1038/ncomms14836

13. Helmli FS, Scherer S. Adaptive shape from focus with an error estimation in light microscopy. In: Proceedings of the 2nd International Symposium on Image and Signal Processing and Analysis (ISPA 2001). Pula, Croatia; 2001. pp. 188–193. doi:10.1109/ISPA.2001.938626

14. Nayar SK, Nakagawa Y. Shape from focus. IEEE Trans Pattern Anal Mach Intell. 1994;16(8):824– 831. doi:10.1109/34.308479

15. Durand F, Hasinoff SW, Freeman WT. Noise-optimal capture for high dynamic range photography. In: Proceedings of the IEEE Conference on Computer Vision and Pattern Recognition (CVPR); 2010 Jun 13–18; San Francisco, CA. p. 553–560. doi:10.1109/CVPR.2010.5540167

16. Peng H. Bioimage informatics: a new area of engineering biology. Bioinformatics. 2008;24(17):1827–1836. doi:10.1093/bioinformatics/btn346

17. Carpenter AE, Jones TR, Lamprecht MR, Clarke C, Kang IH, Friman O, et al. CellProfiler: image analysis software for identifying and quantifying cell phenotypes. Genome Biol. 2006;7(10):R100. doi:10.1186/gb-2006-7-10-r100

18. Macenko M, Niethammer M, Marron JS, Borland D, Woosley JT, Guan X. A method for normalizing histology slides for quantitative analysis. IEEE International Symposium on Biomedical Imaging: From Nano to Macro, Boston, MA, USA. 2009, pp 1107–1110. doi:10.1109/ISBI.2009.5193250

19. Reinhard E, Ashikhmin M, Gooch B, Shirley P. Color transfer between images. IEEE Computer Graphics and Applications. 2001;21(5):34–41. doi:10.1109/38.946629

20. Goodfellow I, Bengio Y, Courville A. Deep Learning. Cambridge (MA): MIT Press; 2016.

21. LeCun Y, Bengio Y, Hinton G. Deep learning. Nature. 2015;521(7553):436-444. doi:10.1038/nature14539

